# Inward tubulation of the plasma membrane expedites membrane exchange and receptor presentation

**DOI:** 10.1101/315606

**Authors:** Long-fang Yao, Li Zhou, Ruilin Zhang, Liang Cai

## Abstract

The plasma membrane is a crucial barrier between the cell and its external environment, and it also enables efficient passage of materials and information. Membrane vesicle trafficking allows precise delivery of materials but is rather inefficient. The mechanism for efficient membrane exchange remains elusive. Here we describe inward tubulation of the plasma membrane (PM tubes) that extends deep into the cytoplasm. These widespread PM tubes elongate along microtubules and are stabilized by actin filaments and cholesterol. PM tubes are preferred sites for connection between the endoplasmic reticulum and the plasma membrane. PM tubes facilitate receptor presentation at the surface of cells, possibly also shortening the distance for transported cargo to reach the external environment.

**In Brief:** A new type of tubular membrane structures was discovered in cells, revealing a shortcut that cells employ to expedite material exchange with their external environment.

**Highlights:**

- Inward tubulation of the plasma membrane (PM tubes), transiently interacts with the Golgi apparatus
- Microtubule side-binding proteins pull PM tubes, while actin filaments and cholesterol stabilize PM tubes
- PM tubes are preferred sites where ER-PM contacts form in response to increased cytoplasmic calcium concentration
- PM tubes are preferred sites for the surface presentation of GLUT1 upon glucose deprivation

## Introduction

All reactions take place in their specific environment, usually within limited space and time. For cells, biological membrane-bound organelles are where all cellular reactions take place. ‘*Omnis membrane e membrane*’, what Günter Blobel addressed in 1999 highlights the principle of membrane biogenesis (Blobel, 2000), as well as the challenge the eukaryotic cells must overcome, that is, to exchange material, including cargos and membranes, precisely and efficiently between organelles.

The benefit of compartmentalized cellular structures allows the cells to have specialized space to execute specialized reactions efficiently, i.e., the nucleus for storage and transmission of genetic information, lysosomes for recycling material, mitochondria and chloroplasts for energy metabolism, endoplasmic reticulum (ER) and Golgi apparatus for protein synthesis and maturation. Most if not all biological reactions happen in membrane-bound organelles, and membranes not only function as barriers between the cell and its external environment but also provide a physical platform for various reactions (Lippincott-Schwartz, 2011; Steinman et al., 1983).

Besides direct interactions, membrane vesicle trafficking is thought to be the only approach to exchange materials between membrane-bound compartments. Certain materials, usually called cargos, are packaged into vesicles, bound by the source membrane, and depart from the source compartment. These vesicles are transported by motor proteins along tracks formed by cytoskeletal filaments. When the cargo-containing vesicles reach the target compartment, the vesicle membrane fuses with the target membrane, and the cargos are released. Simultaneously, the source membrane is being delivered to the target compartment. In eukaryotic cells, material exchange between the compartments is achieved by membrane vesicle trafficking, including transports between organelles, and between the internal and the external environment of the cell. Years of studies in many laboratories around the world have tempted to illustrate detailed mechanisms of almost every aspect of membrane vesicle trafficking (Bonifacino and Glick, 2004; Hutagalung and Novick, 2011; Lippincott-Schwartz et al., 2000). However, the question how to achieve efficient transportation of material, especially membrane, between compartments is still elusive.

Recently, various membrane contact sites have been reported (Chinnapen et al., 2012; Friedman et al., 2011; Giordano et al., 2013; Manford et al., 2012; Rowland et al., 2014; Stefan et al., 2011; Tavassoli et al., 2013). Most of these structures are formed between ER and other membrane-bound organelles, e.g., mitochondria, peroxisomes, and the plasma membrane. These reported structures play important roles in lipid biogenesis (Rowland and Voeltz, 2012), as well as the homeostasis of the contacting organelles. Direct contact between ER and corresponding organelles could achieve efficient material exchange inside the cell. But, how the cell efficiently exchanges material with its external environment?

Here, we describe a tubular membrane structure that widely exists inside cells, extending from the plasma membrane (PM) to the cytosol. We show that these PM tubes are highly dynamic, and have unique kinetic properties. We further determined how these PM tubes expedite membrane exchange between the plasma membrane and contacting organelles, and how they facilitate receptor presentation. Both mechanisms contribute to efficient material exchange between the internal and the external environment of the cell.

## Results

### A widely existing tubular membrane structure inside cells

To visualize membrane dynamics, we labeled the plasma membrane by GFP C-terminally fused with a CAAX sequence from H-Ras (Homo sapiens NP_005334.1, amino acid 170-189). We noticed fluorescent signal from the cell surface as well as some internal structures. Interestingly, besides spheres which we believe were previously reported as trafficking vesicles, GFP-CAAX labeled internal structures also include tubular structures (Figure 1A,1C). These GFP-CAAX labeled tubular membrane structure (tubes for short), are lying underneath the plasma membrane, spanning from one side of a cell to the other side, or from the center to the edge of a cell (Figure 1A-1D). In addition, the tubes were observed in cells with the plasma membrane labeled by fluorescent proteins fused with tetraspanin family proteins (CD9 in Figure 4A).

**Figure 1.**
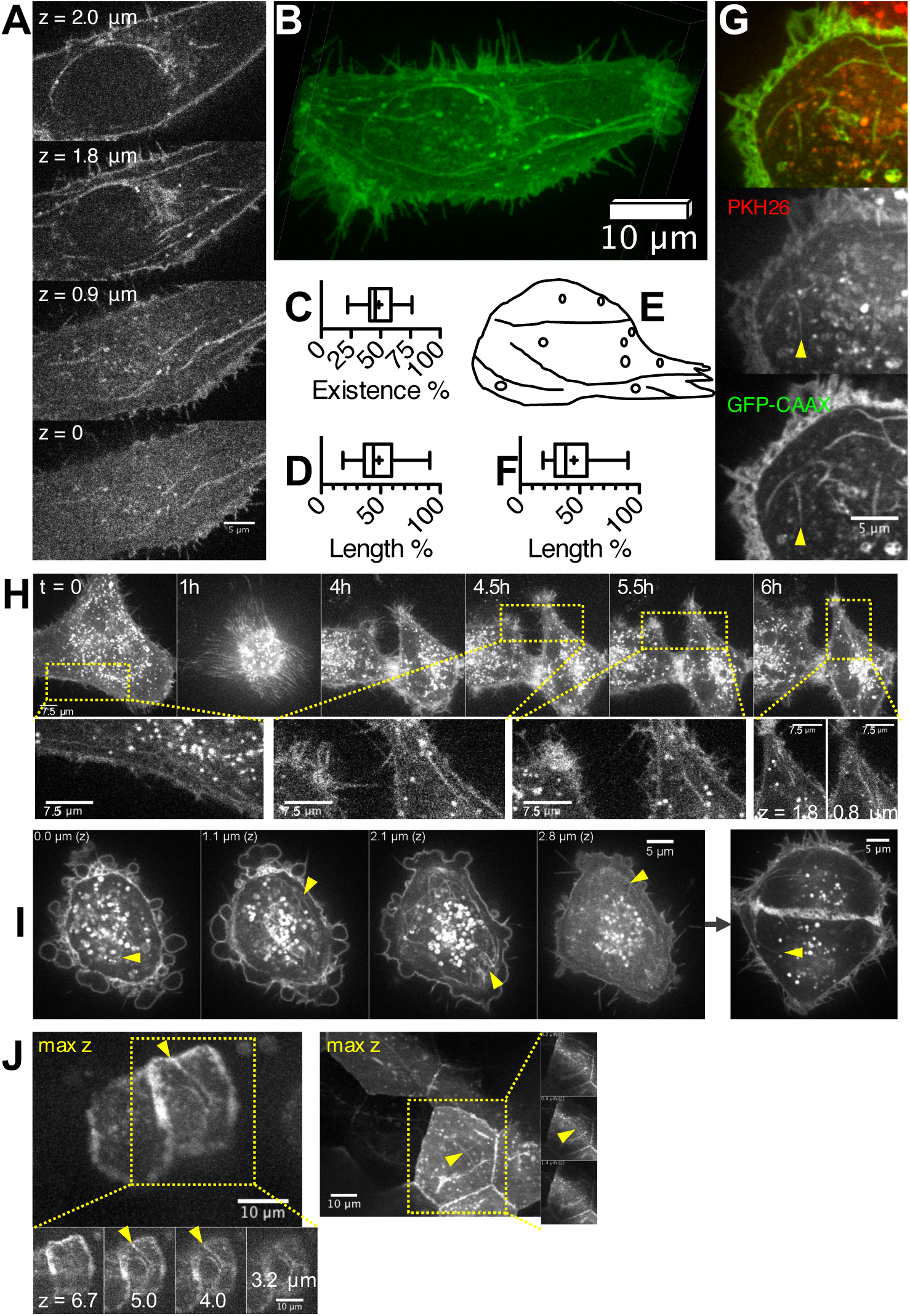
Widely existence of tubular membrane structures. A- HeLa cells expressing GFP-CAAX were lively imaged by spinning disk confocal microscopy (SDCM). Slices of a Z stack from optical sectioning are presented as montage. Scale bar = 5 μm. B- Slices from (A) were imported into Imaris (Bitplane), background subtracted, smoothened using the default Gaussian filter, volume rendered and displayed. C- The percentage of GFP-CAAX expressing HeLa cells with the tubular membrane structures, presented as box and whisker plot (5-95 percentile). D- For cells as in (C), the length of the tube was divided by the long cell axis, and the percentage presented as box and whisker plot. E- Schematic diagram of the tubular membrane structures we discovered (lines), with circles representing membrane vesicles. F- HeLa cells were trypsinized, stained with PKH26 dye, plated on fibronectin coated glass bottom dishes, and lively imaged by SDCM. PKH26 labeled tubular structures were quantified and presented as in (D). G- HeLa cells expressing GFP-CAAX were stained and imaged as in (F). Yellow arrowheads indicate a GFP-CAAX and PKH26 double positive tube. Quantification on maximum intensity projections of Z stacks from optical sectioning shows 97.5 ± 9.3% GFP-CAAX tubes are PKH26 positive (mean ± standard deviation). H- HeLa cells stably expressing GFP-CAAX were imaged by SDCM overnight. The cell underwent cell division. Expanded views of yellow dash lines boxed regions are displayed in the lower panels. I- HeLa cells as in (H), slices of a Z stack from optical sectioning are presented. Yellow arrowheads indicate the GFP-CAAX marked tubes. Gray arrow points to the resulting cells after the division. J- 7 dpf *Tg*(*EF1a*:myr-Tdtomato) zebrafish larva was anesthetized and embedded in low melting agarose in lateral position. Notochord sheath cells were lively imaged by SDCM. Maximum intensity projections of Z stacks are displayed, with expanded view showing individual slices of yellow dash lines boxed regions. Yellow arrowheads indicate the myr-Tdtomato marked tubes.

To ascertain whether the tubes observed are caused by the expression of fluorescent protein fusions, we used live-cell imaging to monitor dye-labeled membrane structures in real time. PKH26 is a widely used lipophilic dye and labels biological membrane by inserting its long aliphatic tails into the lipid regions of the membrane. The length of PKH26 labeled tubular structures had similar distribution as GFP-CAAX labeled tubes (Figure 1F). Additional experiments demonstrated that GFP-CAAX and PKH26 labeled the same tubes (Figure 1G). In summary, using either fluorescent protein fusions or fluorescent lipid, we discovered a novel tubular structure that widely exists inside cells.

We first discovered the tubes in HeLa cells. Then we confirmed that they could be observed in a variety of commonly used cell lines, including 293FT, COS7, NIH/3T3, Raw 264.7, MDCK, NRK and HT-1080 (data not shown). Strikingly, these tubes seemed to be stable through the cell cycle (Figure 1H). When a cell divides, the plasma membrane undergoes dramatic reformation. However, the tubes persist and exist in dividing cells (Figure 1I).

Limited literature searching suggests that at least in developing worm embryo (8-cell stage, Chen et al., 2014) and zebrafish larva (notochord sheath cells, 7-day post fertilization, Dale and Topczewski, 2011), there are tubular membrane structures in animals expressing fluorescent protein at the plasma membrane. We confirmed the second in Figure 1J.

Recent advances in super-resolution microscopy have substantially increased the spatial resolution. We then performed live-cell analysis using a super-resolution microscopy, Airyscan from ZEISS (Figure 2A). The diameter of the tubes (labeled by mMaple3-CAAX) was measured to be 253 ± 49 nm (mean ± standard deviation, Figure 2E). Both ends of a tube spanning across a cell are shown, with one end residing closely to the plasma membrane (Figure 2B).

**Figure 2.**
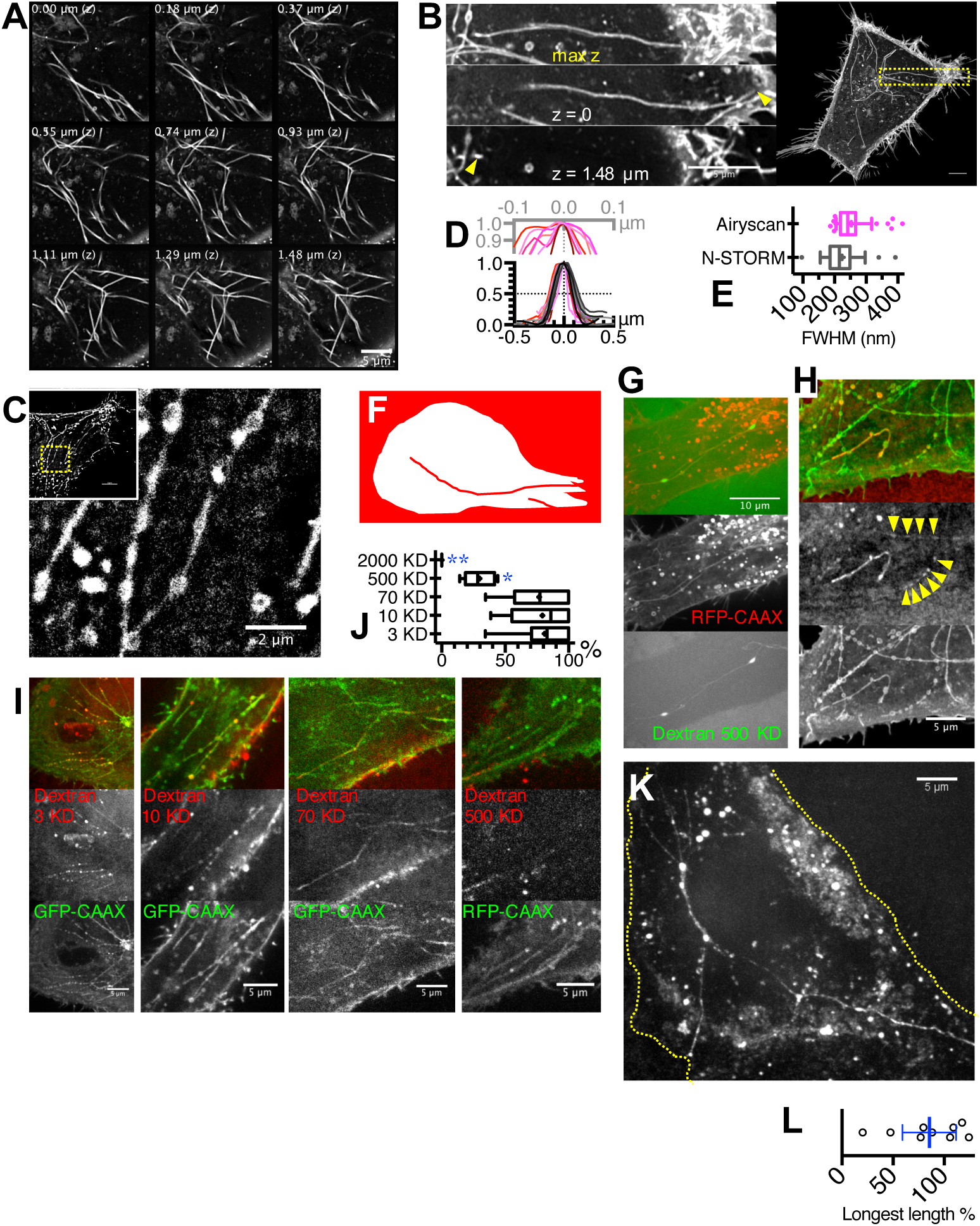
Inward tubulation of the plasma membrane. A- HeLa cells expressing GFP-CAAX were lively imaged by the LSM 880 laser scanning microscope with Airyscan (ZEISS). Slices of a Z stack from optical sectioning are presented as montage. Scale bar = 5 μm. B- Maximum intensity projection of an Airyscan Z stack from (A) is displayed on the left, with expanded view of yellow dash line boxed region on the right. Two slices of the Z stack presented with yellow arrowheads indicating the two ends of the tube. C- HeLa cells expressing mMaple3-CAAX were fixed with 4% paraformaldehyde and 0.5% glutaraldehyde, and imaged by a N-STORM super-resolution microscope (Nikon). Yellow dash line boxed region of the cell is displayed in expanded view. D- Pixel intensities around the tubes were normalized and plotted based on their distances to the axis. A detailed view of the distribution within 0.1 μm of the axis is displayed in the upper panel. Grey lines were from 7 Airyscan captured tubes and pink lines were from 7 N-STORM captured tubes. E- Tube diameters were measured by the full-width-half-maximum (FWHM) method, and presented as box and whisker plots (10-90 percentile) with the outliners. F- Schematic diagram of dextran loading assay: the cells were plated on fibronectin coated glass for 12 hr, and incubated with lysine-fixable fluorescent dextran containing culture media. G- HeLa cells expressing GFP-CAAX were treated as in (F) with 500,000 MW dextran for 5 min, and lively imaged by SDCM. H- HeLa cells expressing GFP-CAAX were treated as in (F) with 70,000 MW dextran for 10 sec, fixed and imaged by Airyscan. Yellow arrowheads indicate dextran containing membrane bubbles. I- Fluorescent dextran with different molecular weights were used (KD, kilodalton). HeLa cells expressing GFP-CAAX were treated as in (F) for 10 sec, fixed and imaged by SDCM. J- For cells as in (I), the length of the dextran loaded region was divided by the length of the tube, and the percentage presented as box and whisker plots. * indicates that the cells were incubated with 500,000 MW dextran for 1 min, while ** were incubated with 2,000,000 MW dextran for 3 min. K- Plain HeLa cells were treated as in (F) with 70,000 MW dextran for 10 sec, fixed, imaged by SDLM, and presented as maximum intensity projection of a Z stack from optical sectioning. L- For cells as in (K), the length of the longest tube loaded with dextran was divided by the long cell axis, and the percentage presented as scatter dot plot.

Using a combination of paraformaldehyde and glutaraldehyde reagents, we were able to fix the tubes for N-STORM super-resolution imaging (Figure 2C). We measured the diameter of the tubes (labeled by mMaple3-CAAX) to be 224 ± 60 nm (mean ± standard deviation, Figure 2E). With N-STORM, we were able to observe the hollow structure of the tubes in some cross sections (Figure 2D). Notably, previously smooth tubes bulged to form a lot of membrane bubbles. We think that bulging may be due to an un-synchronized fixation of the fast flowing tubular membrane.

### Tubular membrane structures have openings on the plasma membrane

Inspired by the hollow structure, we were curious whether we could mark the tubes by what they transport. Dextran is a complex branched polysaccharide, taken by cells via endocytosis, and is used as a classical marker for membrane vesicle trafficking. We added fluorescent dextran (10,000 MW) into the cell culture media and pulsed for 15 min, then chased with plain culture media and imaged the living cells. Consistent with previous knowledge that dextran is taken and transported as vesicles, we observed that most fluorescent signals were vesicles of various sizes (data not shown). Moreover, we were able to find tubular structures with bright fluorescence. By using a fluorescent dextran with a large molecule weight (500,000 MW) and reducing the pulse time to 5 min, most of the dextran signals were limited to tubular structures (Figure 2G). Furthermore, fluorescent dextran also marked the tubes labeled by RFP-CAAX. Thus, we show that the tubes inside cells with the hollow structure may participate in dextran transport.

After careful examination of fluorescent dextran marked tubes, we noticed several interesting features. Only some portions of GFP-CAAX labeled tubes were marked with fluorescent dextran, while all fluorescent dextran marked tubular structures were labeled by GFP-CAAX. Every dextran marked tube had at least one end very close to the plasma membrane. Furthermore, the end close to the plasma membrane always had a stronger fluorescent signal compared with the other end. Based on these features, we hypothesized that dextran was siphoned into the tube due to its opening on the plasma membrane.

To test our hypothesis, we added fluorescent dextran (70,000 MW) into the cell culture media for only 10 sec before fixation. Strikingly, fluorescent dextran siphoned in and vividly marked tubular structures spanning across the cell (Figure 2K,2L), with enriched signals from fixation generated bulged membrane bubbles (Figure 2H).

We measured the average diameter of the tubes to be 224~253 nm, and thus dextran with a comparable diameter would not be able to siphon in and mark the tubes. We incubated cells with different sized fluorescent dextran (10,000 MW, 30,000 MW, 70,000 MW, 500,000 MW, 2,000,000 MW), and measured the labeling efficiency (Figure 2I,2J). Because the concentration of fluorescent dextran we used was 2 mg/ml, the change of fluid viscosity upon the addition of dextran was minimal and could be neglected. While dextran with a molecule weight less than 70,000 siphoned into the tubes and marked over 80% length of the structures, dextran 2,000,000 MW could not mark any internal cellular structure. Respectively, the Stokes radius of 57 KD and 157 KD dextran are 5.5 nm and 9.1 nm (Peters, 1984). Together, we showed that these widely existing tubes have openings on the plasma membrane.

### Inward tubulation of the plasma membrane

To confirm a direct connection between the plasma membrane and the tubes, we photo-converted a portion of the membrane from green fluorescence to red in cells expressing mMaple3-CAAX (Wang et al., 2014), and performed live-cell imaging of converted membrane. A red tube initiated from a region with no visible vesicle or organelle, and protruded into a non-illuminated region (Figure 3G), suggesting that the tubes were built using the material from the initiation plasma membrane. This is consistent with the observation that the tubes have openings on the plasma membrane. In addition, upon green-to-red conversion, previously existing tubes immediately obtain red signals, indicating a fast membrane flow on the tubes.

**Figure 3.**
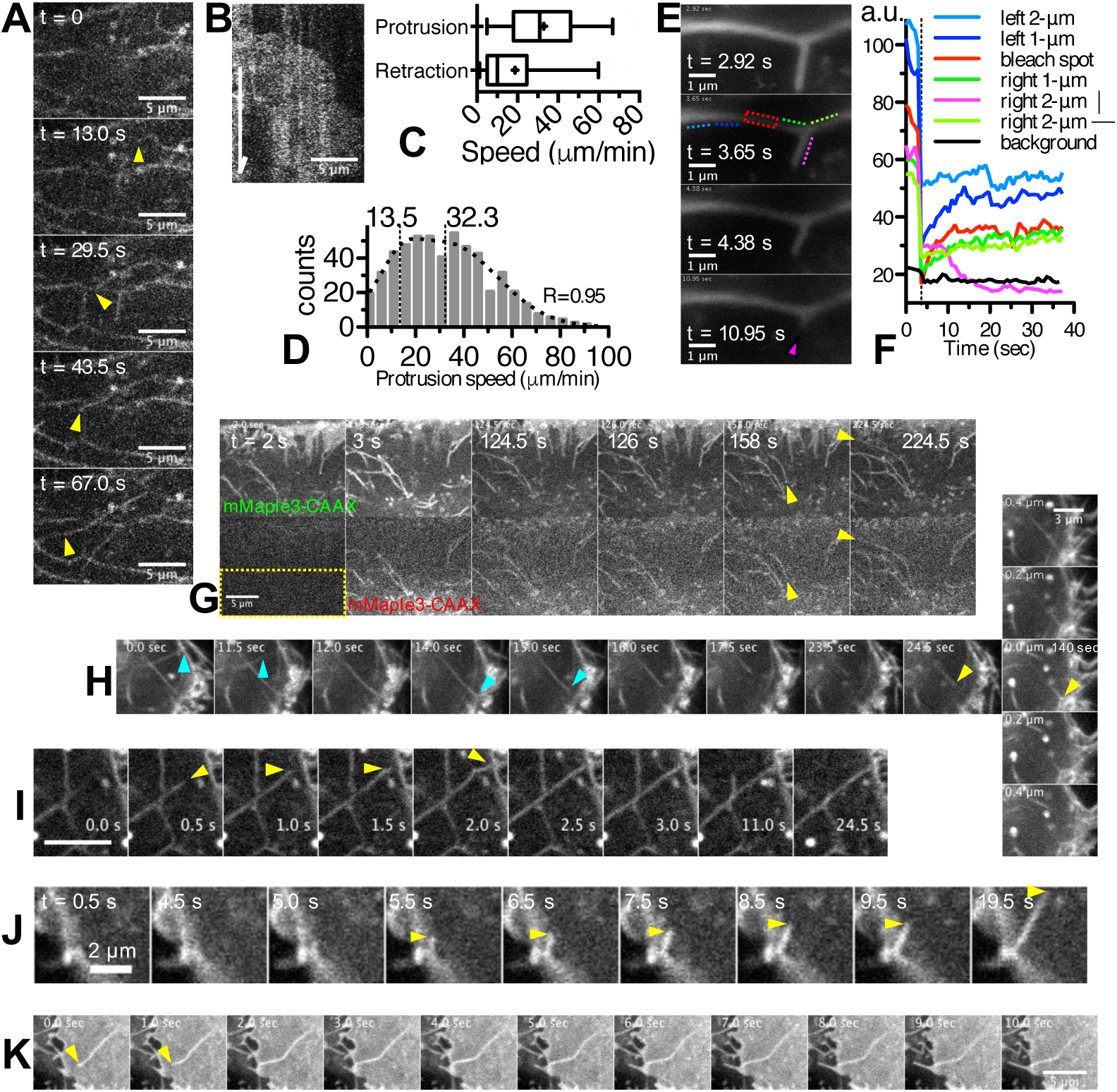
Dynamics of the plasma membrane tubes. A- HeLa cells expressing GFP-CAAX were lively imaged by SDCM and presented as timelapse montage. Yellow arrowheads indicate one end of a protruding tube. Scale bar = 5 μm. B- Kymograph of the protruding tube in (A). Horizontal line = 5 μm; vertical line = 1 min. C- For cells as in (A), kymograph analysis as in (B) was used to calculate the dynamic parameters, which were presented as box and whisker plots. D- The protrusion speed of (C) presented in a histogram. Sum of two Gaussians was fit to the data to calculate the means. E,F- HeLa cells expressing GFP-CAAX were lively imaged by a laser scanning microscope. The red dash line boxed region was photobleached. Pixel intensities of different regions of the tube were plotted over time. Pink arrowhead indicates the retracting tube. G- HeLa cells expressing mMaple3-CAAX were lively imaged by SDCM. Yellow dash line boxed region was illuminated with 405 nm laser for 1 sec to convert some green fluorescent mMaple3-CAAX to red fluorescent mMaple3-CAAX. Images are presented as timelapse montage. Yellow arrowheads indicate a tube that imitates from the illuminated region and protrudes into the non-illuminated region. H,I,J,K- HeLa cells expressing GFP-CAAX were lively imaged by SDCM and presented as timelapse montage horizontally. In (H), slices of the Z stack from optical sectioning at t = 140 sec are presented as montage vertically. Cyan arrowheads indicate one end of a retracting tube, and yellow arrowheads indicate ends of protruding tubes.

To measure the directionality of the tubular membrane flow, we performed fluorescence recovery after photo-bleaching (FRAP) analysis (Figure 3E-F). Due to the fast membrane flow, fluorescence of all measured regions decreased upon bleaching. During the recovery phase, within a limited time, only one side of the bleached region showed increased fluorescence. The estimated membrane flow speed (2~3 μm in 6 sec) is consistent with calculated fast protrusion speed (32.3 μm/min, Figure 3D) of the tubes. Thus, we conclude that the membrane flow on the tubes is unidirectional.

Careful examination of live-cell imaging data revealed further evidence supporting the direct connection between the plasma membrane and the tubes. There are tubes retracting from the plasma membrane (Figure 3H). There are tubes protruding and connecting to the plasma membrane (Figure 3H,3I). And tubes initiating from the plasma membrane (Figure 3J). Infrequently, there were tubes connecting to the plasma membrane and supporting filopodia outgrowth (Figure 3K). Together, we discovered that inward tubulation of the plasma membrane forms tubes spanning across the cell.

### Plasma membrane tubes are not tubular mitochondria or tubular endoplasmic reticulum

Mitochondria, found in most eukaryotic cells, play an important role in energy production. Recent data indicate that besides fission and fusion, dynamic tubulation of mitochondria is an essential mechanism for its network formation (Wang et al., 2015). To clarify that the plasma membrane (PM) tubes are not tubular mitochondria, we labeled mitochondria with a fluorescent protein (FP) fused with the C terminus of ActA, and labeled PM tubes with FP-CAAX. With FP swapping combination, in neither case, the tubes colocalize with mitochondria (Figure 4C).

**Figure 4.**
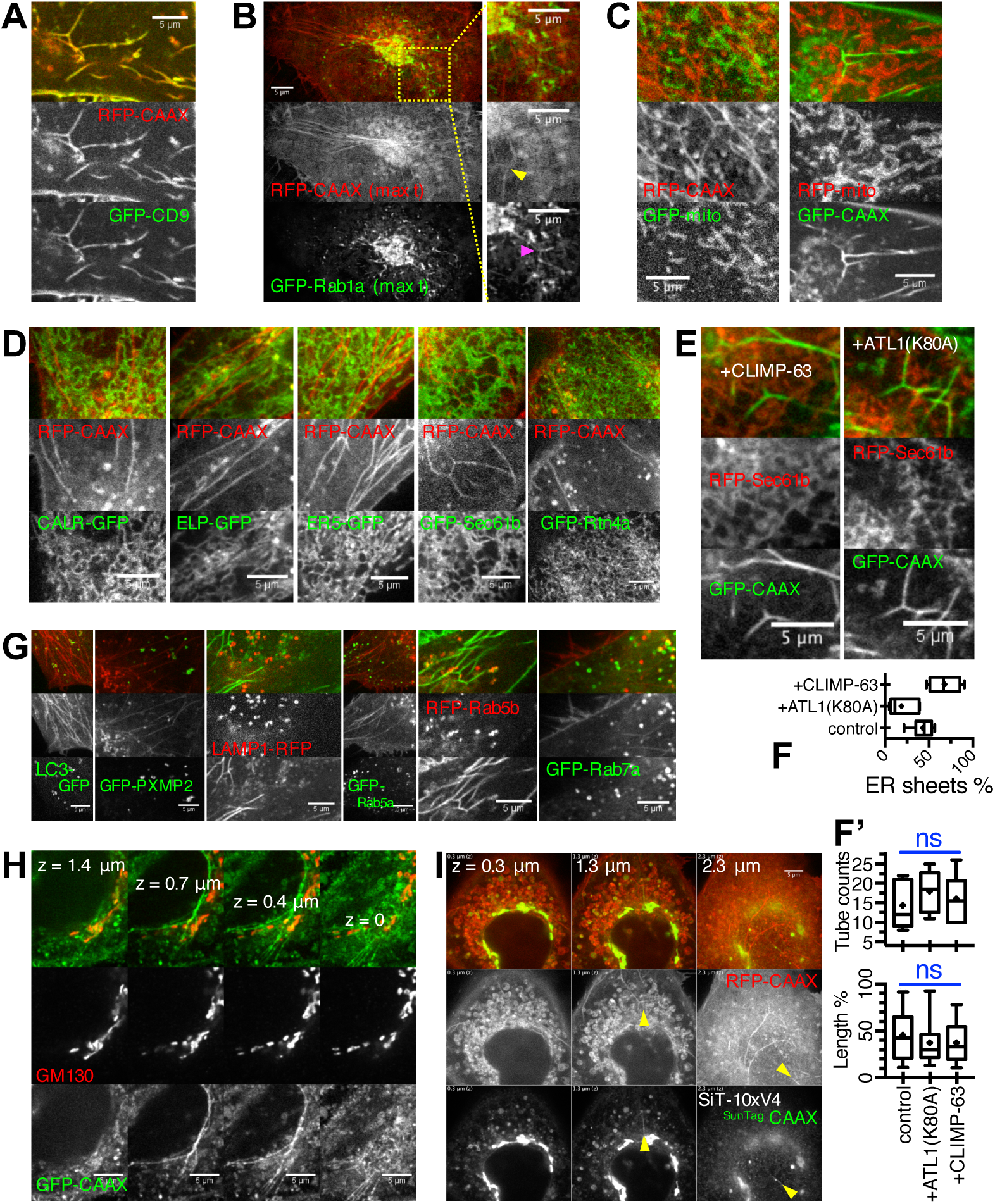
The plasma membrane tubes are uncharacterized membrane structures. A- HeLa cells expressing RFP-CAAX and GFP-CD9 were lively imaged by SDCM. Scale bar = 5 μm. B- HeLa cells expressing RFP-CAAX and GFP-Rab1a were lively imaged by SDCM. Maximum intensity projection of a timelapse is displayed on the left, with expanded view of yellow dash line boxed region on the right. Pink arrowhead indicates a protruding GFP-Rab1 positive tube, and yellow arrowhead indicates a protruding RFP-CAAX positive tube. C- HeLa cells expressing markers for the plasma membrane and mitochondria (mito) were lively imaged by SDCM. D- HeLa cells expressing markers for the plasma membrane and endoplasmic reticulum (ER) were lively imaged by SDCM. Five different markers for the endoplasmic reticulum were used: calreticulin (CALR), ERD-2-like protein 1 (ELP), GFP fusion with KDEL sequence (ER5), Sec61b and reticulon 4a (Rtn4a). E- HeLa cells expressing GFP-CAAX and RFP-Sec61b were co-transfected with either atlastin-1(K80A) or CLIMP-63, and lively imaged by SDCM. F,F’- For cells as in (E), the length of the tube was divided by the long cell axis, the area of ER sheets was divided by the area of the cell, or the number of tubes in the cell was counted, and the percentage presented as box and whisker plots. One-way ANOVA reports the difference is not significant (ns). G- HeLa cells expressing markers for the plasma membrane and known organelles were lively imaged by SDCM. Six different markers for five known organelles were used: autophagosome (LC3), peroxisome (PXMP2), lysosome (LAMP1), early endosome (Rab5a, Rab5b) and late endosome (Rab7a). H- HeLa cells expressing GFP-CAAX were fixed, stained with antibody against a *cis*-Golgi matrix protein (GM130), and imaged by SDCM. Slices of a Z stack from optical sectioning are presented as montage. I- SiT, a marker for the trans-Golgi network, was fused with 10xV4, co-expressed with ^SunTag^CAAX in HeLa cells, and lively imaged by SDCM. Slices of a Z stack from optical sectioning are presented as montage. Yellow arrowheads indicate the double positive tubes.

The endoplasmic reticulum (ER) is critical for both protein and lipid synthesis in eukaryotic cells. ER is a network of tubes and sheets (Voeltz and Prinz, 2007; Voeltz et al., 2006). To clarify that the PM tubes are not tubular ER, we labeled ER with GFP fused with various ER residing proteins (calreticulin, ERD-2-like protein 1, Sec61b, reticulon 4a) or a KDEL sequence. In none of the cases, the PM tubes overlapped with any ER structure (Figure 4D). Even though the tubular ER spread, extended and covered almost all the cytoplasmic space, PM tubes always stayed out of the ER territory.

To further confirm that there is no interaction between PM tubes and ER, we shifted the balance between tubular ER and ER sheets. As previously reported, when CLIMP-63 was over-expressed in cells, the percentage of tubular ER decreased (Klopfenstein et al., 2001), while overexpressing ATL1(K80A) increased long and unbranched ER tubules (Hu et al., 2009), (Figure 4E-F). In both cases, neither the length of PM tubes nor the number of tubes in the cell changed (Figure 4F’). Together, we conclude that PM tubes are not tubular ER nor mitochondria.

### Plasma membrane tubes transiently interact with the Golgi apparatus

We showed that the plasma membrane inward tubulated into the cell, and maintained openings on the plasma membrane. We were curious where the other ends of these tubes reside. Using a panel of well-characterized markers for intracellular organelles, we performed colocalization analysis. Although small portions of Rab5a/Rab5b positive early endosomes aligned along the tunnels, we failed to detect any connection between the ends of PM tubes with autophagosomes, peroxisomes, lysosomes, early or late endosomes (Figure 4G).

The Golgi apparatus processes and packages cargos into vesicles before delivering them to their targets. If the tubes involved in delivering cargos, it is likely that they cease at the Golgi apparatus. We labeled the Golgi apparatus either by expressing GFP fused with Golgi residing proteins (β1,4-galactosidase, sialyltransferase; data not shown), or by immunofluorescence using an antibody against GM130 (*cis*-Golgi marker). In cells with both the tubes and the Golgi labeled, the intracellular ends of the tubes somewhat overlapped with the Golgi. However, using dual-color live-cell imaging, we could not ascertain whether or not the tubes protruding from the plasma membrane interact with the Golgi apparatus.

To confirm the potentially transient interaction between the Golgi and PM tubes, we employed the SunTag fluorescence tagging strategy (Tanenbaum et al., 2014; Yao et al., 2017). In cells only expressing GCN4-sfGFP-GB1-CAAX (^SunTag^CAAX, GCN4 is a single-chain variable fragment antibody against V4 peptide), both the plasma membrane and the tubes, as well as trafficking vesicles, were labeled by GFP-CAAX. In control cells expressing GCN4-sfGFP-GB1 (^SunTag^cyto) and an array of 10 copies of V4 peptide fused with sialyltransferase (SiT-10xV4), fluorescence was cytoplasmic and no membranous structure was labeled (data not shown), consistent with the finding that 10xV4 part of SiT-10xV4 is non-cytoplasmic. Interestingly, in cells expressing ^SunTag^CAAX and SiT-10xV4, neither the plasma membrane nor most of the tubes were labeled, but the Golgi was labeled. It seemed that the ^SunTag^CAAX signal was sequestered in the Golgi apparatus. Notably, the SunTag part of ^SunTag^CAAX is cytoplasmic as of ^SunTag^cyto, with the difference that ^SunTag^CAAX resides on the plasma membrane, or on the cytosolic surface of ER and Golgi during its maturation (Apolloni et al., 2000; Choy et al., 1999). These data suggested that the plasma membrane continuously has transient interaction with the Golgi apparatus through the tubes, and this interaction provides membrane continuity and must surpass the topological barrier.

Post-Golgi carriers are important components of intracellular transport, and some of the post-Golgi carriers are membrane tubules being pulled out of the *trans*-Golgi network (Almeida et al., 2011; Kreitzer et al., 2003; Luini et al., 2008; Russo et al., 2016). To ascertain that the PM tubes are not post-Golgi tubules, we used dual-color live-cell imaging to compare the dynamics of Rab1a labeled post-Golgi tubules, and RFP-CAAX labeled PM tubes in real time. In summary, although the PM tubes transiently interact with the Golgi, they are not post-Golgi tubules.

### Actin filaments stabilize plasma membrane tubes

PM tubes are dynamic membrane structures, protruding and retracting all the time (Figure 3C). Other known intercellular structures with analogical dynamics are actin filaments and microtubules (Fletcher and Mullins, 2010). We postulated that some cytoskeletal filaments play important roles in regulating the dynamics of the tubes.

First, we labeled the actin filaments with LifeAct-RFP and performed colocalization analysis with GFP-CAAX labeled PM tubes. Although spatially some actin filaments extensively colocalize with some PM tubes (Figure 5B), their colocalization has a temporal difference. In protrusions, PM tubes always precede actin filaments elongation (Figure 5A,5C-D). In retractions, the actin filaments have to disappear prior to the tubes (Figure 5C-D).

**Figure 5.**
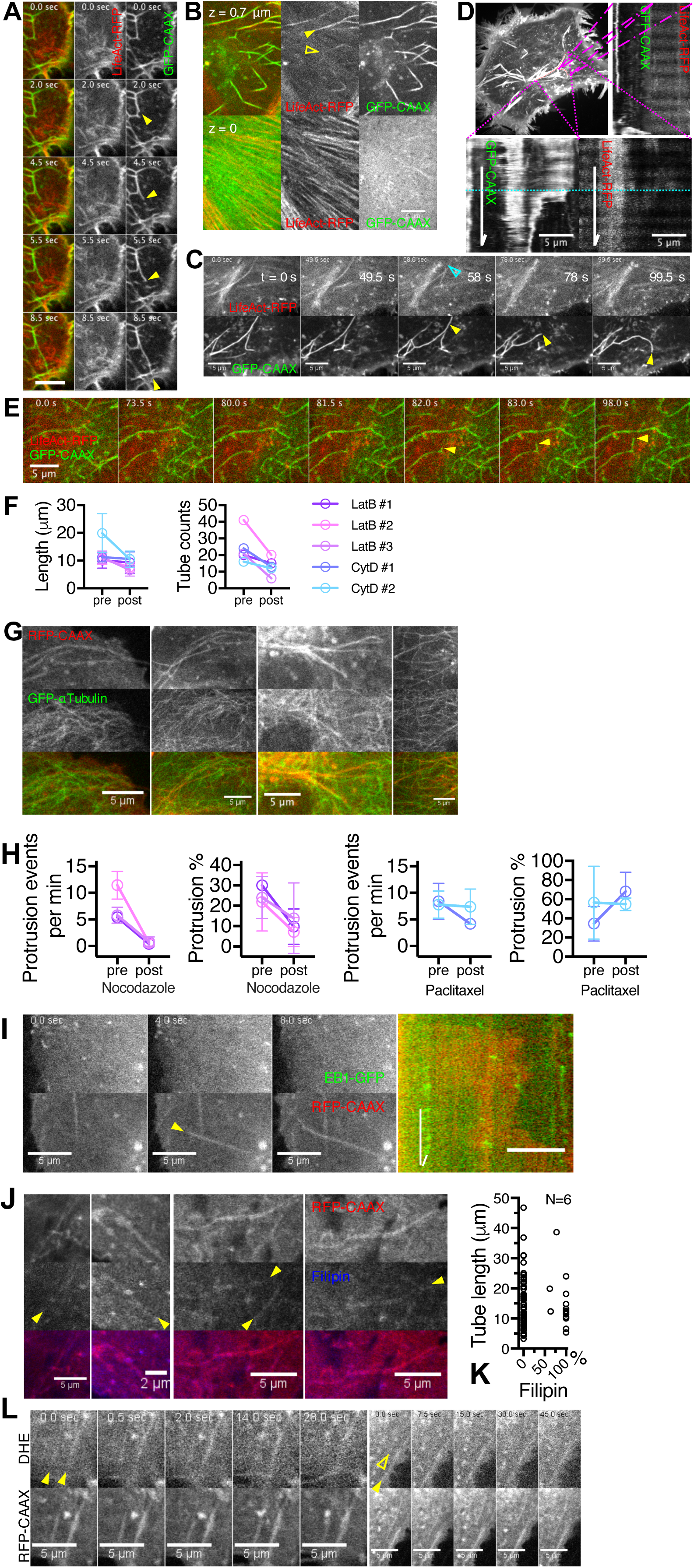
Regulation of the dynamics of the plasma membrane tubes. A- HeLa cells expressing GFP-CAAX and LifeAct-RFP were lively imaged by SDCM and presented as timelapse montage. Yellow arrowheads indicate one end of a protruding tube. Scale bar = 5 μm. B- For cells as in (A), slices of a Z stack from optical sectioning are presented as montage. Yellow arrowhead indicates a double positive tube, and open arrowhead indicates a tube without LifeAct-RFP fluorescence. C- For cells as in (A), timelapse images are presented as montage. Yellow arrowheads indicate a tube retracts, turns, and protrudes. Open arrowhead indicates a disassembling actin filament. D- For the cell in (C), maximum intensity projection of its timelapse is displayed on the up left corner, with kymographs from the pink lines displayed on the right and in the lower panels. The cyan dash line indicates the time when the actin filament disassembled but the tube had not retracted. Horizontal lines = 5 μm; vertical lines = 1 min. E- HeLa cells as in (A) were treated with 20 μM latrunculin B for 30 min, imaged by SDCM and presented as timelapse montage. F- For cells as in (A), the length of the tube and the number of tubes in the cell, before and after 20 μM latrunculin B (LatB) or 200 nM cytochalasin D (CytD) 30-min treatment, was measured or counted, grouped by the experiments and presented as mean ± 95% confidence intervals. G- For cells expressing RFP-CAAX and GFP-αTubulin were lively imaged by SDCM and regions from four different cells are displayed. H- For cells as in (G), the number of protruding tubes and its percentage of the total number of tubes, before and after 10 μM nocodazole or 2 μM paclitaxel 30-min treatment, was counted or calculated, grouped by the experiments and presented as mean ± 95% confidence intervals. I- HeLa cells expressing RFP-CAAX and EB1-GFP were lively imaged by SDCM and presented as timelapse montage. Yellow arrowhead indicates one end of a protruding tube. Kymograph of the protruding tube is displayed on the right. Horizontal line = 5 μm; vertical line = 30 sec. J- HeLa cells expressing RFP-CAAX were fixed and stained with filipin. Regions from four different cells are displayed. Yellow arrowheads indicate some double positive tubes. K- For cells as in (J), the length of RFP-CAAX positive tubes and their filipin stained percentage were presented as scatter plot. L- HeLa cells expressing RFP-CAAX were incubated with dehydroergosterol (DHE), lively imaged by SDCM and presented as timelapse montage. Regions from two different cells are displayed. Yellow arrowheads indicate double positive tubes, and open arrowhead indicates a dynamic tube without DHE labeling.

Second, we used a panel of small molecules inhibitors to perturb the actin dynamics in the cells. When the actin assembly was inhibited by latrunculin B treatment (Figure 5E), PM tubes still protruded, suggesting that the actin filaments do not initiate the elongation of the tubes. In addition, either cytochalasin D or latrunculin B treatment reduced the length of the tubes and the number of PM tubes in the cell. Thus, actin assembly does not initiate but stabilize PM tubes.

### Microtubule dynamics drives plasma membrane tubes

To test the role of microtubules in regulating the dynamics of PM tubes, we performed colocalization analysis with RFP-CAAX labeled tubes and α-tubulin-GFP labeled microtubules (Figure 5G). Colocalization of both fluorescent signals suggested a strong connection between PM tubes and microtubules.

Nocodazole and paclitaxel are two chemical inhibitors that affect microtubule dynamics differently. While nocodazole interferes the assembly and accelerates the disassembly of microtubules, paclitaxel stabilizes the existing filaments and promotes microtubule assembly. In cells treated with nocodazole for 30 min, microtubules were completely disassembled, and the number of protruding PM tubes significantly decreased. However, the tubes persist (Figure 5H). It is consistent with previous observations that actin filaments stabilize the tubes.

In cells treated with paclitaxel for 30 min, we still observed rapid PM tube elongation suggesting microtubule polymerization is not the driving mechanism for PM tubulation, although calculated microtubule polymerization speed is 30 μm/min (Parsons and Salmon, 1997), a close match to our measured tube fast protrusion speed (32.3 μm/min).

To confirm the elongation of PM tubes does not interact with the plus ends of microtubules, we labeled the plus ends of microtubules with EB1-GFP and examined where the ends of RFP-CAAX labeled PM tubes reside. The plus ends of microtubules were observed as comets, and these EB1-GFP positive comets move quickly from the center to the periphery. However, the ends of the PM tubes didn’t overlap with any EB1-GFP signal (Figure 5I). Besides, it is difficult to image how inward tubulated plasma membrane interacts with known microtubule tip attachment complexes (Allan and Vale, 1994; Allan and Vale, 1991; Waterman-Storer et al., 1995). Together, our results indicate that PM tubes elongate along microtubules, by interacting with microtubule side-binding proteins.

### Non-dynamic plasma membrane tubes are enriched with cholesterol

We were curious how the PM tubes remain their shape without stabilizing actin filaments. Cholesterol is a lipid essential for cell viability that plays a critical role in maintaining membrane integrity and is always found in rigid membrane subdomains (Simons and Toomre, 2000). We treated cells with methyl-beta-cyclodextrin (MβCD) to deplete plasma membrane cholesterol. Interestingly, in the cells depleted of cholesterol, the PM tubes were very similar to the ones after cytochalasin D, or latrunculin B treatment (data not shown), suggesting cholesterol stabilizes PM tubes.

Then, we fixed and stained cells with filipin, a fluorescent compound that specifically binds to cholesterol and is widely used as a histochemical marker for cholesterol. We observed labeling of filipin on some PM tubes (Figure 5J-K). Only small portions of PM tubes have partial filipin labeling, suggesting loading cholesterol to a PM tube is a rapid process.

Furthermore, we performed live-cell imaging of cells incubated with a fluorescent cholesterol analog, dehydroergosterol (DHE). We found that the PM tubes were labeled by DHE (Figure 5L), and those non-dynamic tubes had strong DHE labeling. High concentration of cholesterol in membrane lipid results in reduced membrane fluidity. Our results demonstrated that cholesterol plays a role in stabilizing the PM tubes.

### Plasma membrane tubes expedite membrane exchange with the endoplasmic reticulum

Although we have shown that PM tubes are not tubular ER, we wondered whether these two different membrane structures interact with each other. Recent data indicate that extended synaptotagmin protein 1 (E-Syt1), an ER protein, bridges ER and PM upon increased cytoplasmic calcium concentration (Giordano et al., 2013). These ER-PM contact sites were suggested to facilitate direct lipid transfer between membranes (Fernandez-Busnadiego et al., 2015; Schauder et al., 2014). We performed live-cell imaging of cells expressing RFP-CAAX and GFP–E-Syt1. In unstimulated cells, E-Syt1 was exclusively located on the ER network. Then, we treated cells with thapsigargin (Tg), a non-competitive inhibitor of the ER calcium-ATPase pump, to induce calcium release from the ER. Strikingly, with the increase of cytoplasmic calcium, E-Syt1 relocalized to the PM tubes and brought the ER to be in close contact with the PM tubes. 20-sec Tg treatment was sufficient to induce ER-PM contact sites, preferably on PM tubes (Figure 6A-B,6H), without affecting ER morphology (Figure 6C-D). Similarly, some E-Syt3 marked ER-PM contact sites formed on the PM tubes (Figure 6E,6H). In summary, the E-Syt1 and E-Syt3 marked ER-PM contact sites preferably formed on the PM tubes, which were tubular structures extending from the plasma membrane.

**Figure 6.**
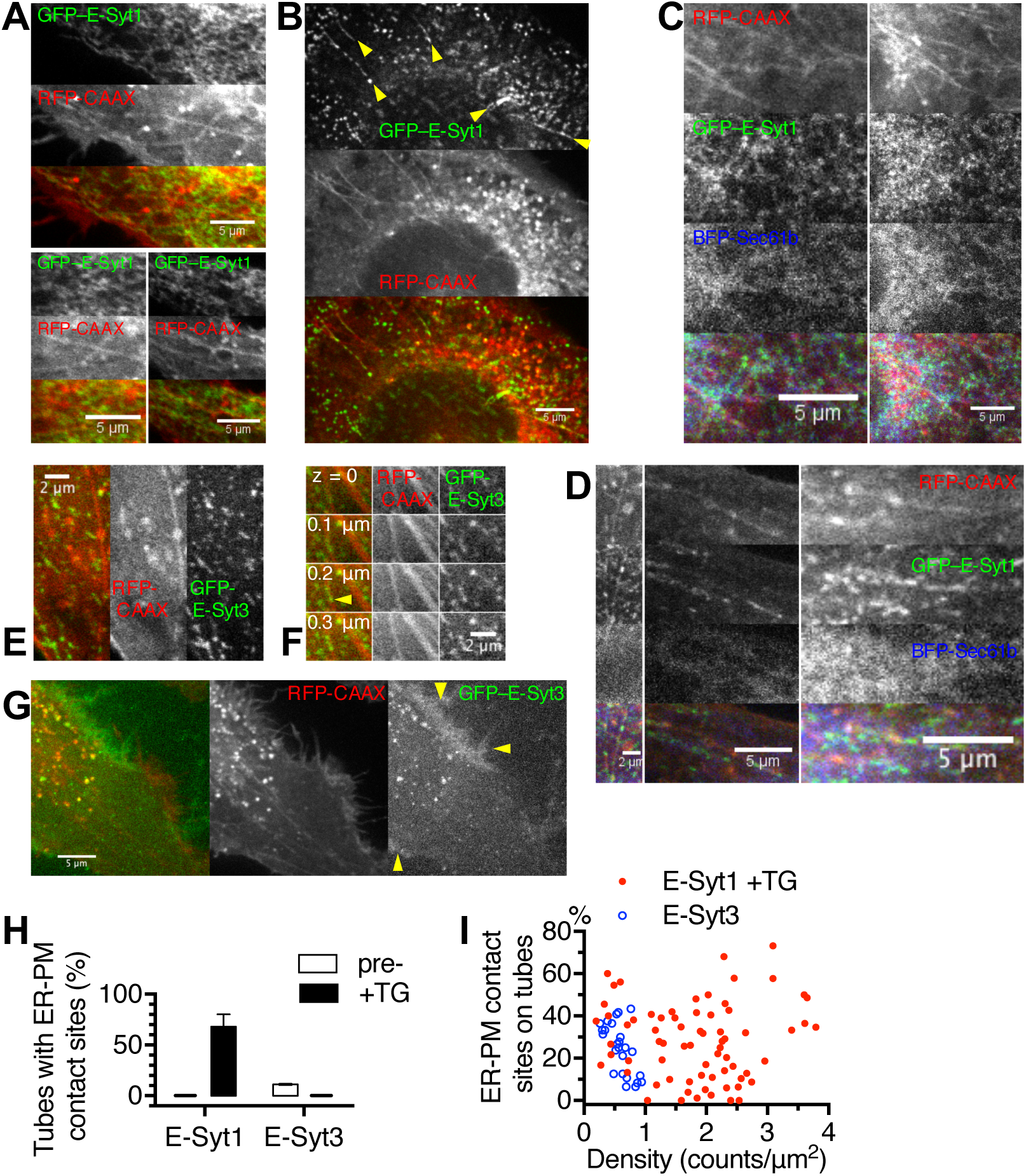
The plasma membrane tubes are the preferred sites to connect the endoplasmic reticulum to the plasma membrane. A- HeLa cells expressing RFP-CAAX and GFP–E-Syt1 were fixed and imaged by SDCM. Regions from three different cells are displayed. Scale bar = 5 μm. B- HeLa cells as in (A) were treated with 200 nM thapsigargin (TG) for 20 sec, fixed and imaged by SDCM. Yellow arrowheads indicate GFP–E-Syt1 dots enriched on tubes. C- HeLa cells expressing RFP-CAAX, GFP–E-Syt1 and EBFP2-Sec61b were fixed and imaged by SDCM. Regions from two different cells are displayed. D- HeLa cells as in (C) were treated with 200 nM thapsigargin (TG) for 20 sec, fixed and imaged by SDCM. Regions from three different cells are displayed. E- HeLa cells expressing RFP-CAAX and GFP–E-Syt3 were fixed and imaged by SDCM. F- For cells as in (E), slices of a Z stack from optical sectioning are presented as montage. Yellow arrowhead indicates a GFP–E-Syt3 dot colocalizes with a tube. G- HeLa cells as in (E) were treated with 200 nM thapsigargin (TG) for 20 sec, fixed and imaged by SDCM. Yellow arrowheads indicate spilled green fluorescence out of the cells. H- For cells in (A-G), the percentage of tubes enriched with ER-PM contact sites, marked by either GFP–E-Syt1 or GFP–E-Syt3, were presented as mean ± standard error of the mean. I- For cells in (H), the density of total ER-PM contact sites in randomly selected regions and the percentage of those ER-PM contact sites colocalizing with tubes were presented as scatter plot.

### Plasma membrane tubes facilitate the surface presentation of receptors

Enlightened by the shortcut that the cell has taken to form the ER-PM contact sites, we wondered whether the PM tubes were the preferred plasma membrane subdomains where receptors were being targeted for presentation.

Glucose is the optimal fuel for most living cells. Uptake of glucose into mammalian cells is mediated mainly by the glucose transporters as the biological membrane is impermeable to glucose. GLUT1 was the first cloned glucose transporter (Mueckler et al., 1985), and widely exists in all types of cells. Immunolocalization of endogenous proteins reveals that GLUT1 dots spread over the plasma membrane (Figure 7A). To investigate the role of PM tubes in GLUT1 surface presentation, we cultured the cells in glucose deprivation conditions for different periods of time. Immediately after replenishing glucose, we fixed, immunostained and examined the localization of endogenous GLUT1. As expected, under glucose deprivation, increased GLUT1 is required on the plasma membrane. These on-demand GLUT1 first enriched on PM tubes and then spread over the plasma membrane (Figure 7B-D). Similarly, after replenishing glucose, the rapid clearing of GLUT1 from the PM tubes preceded the decrease of GLUT1 from the plasma membrane.

**Figure 7.**
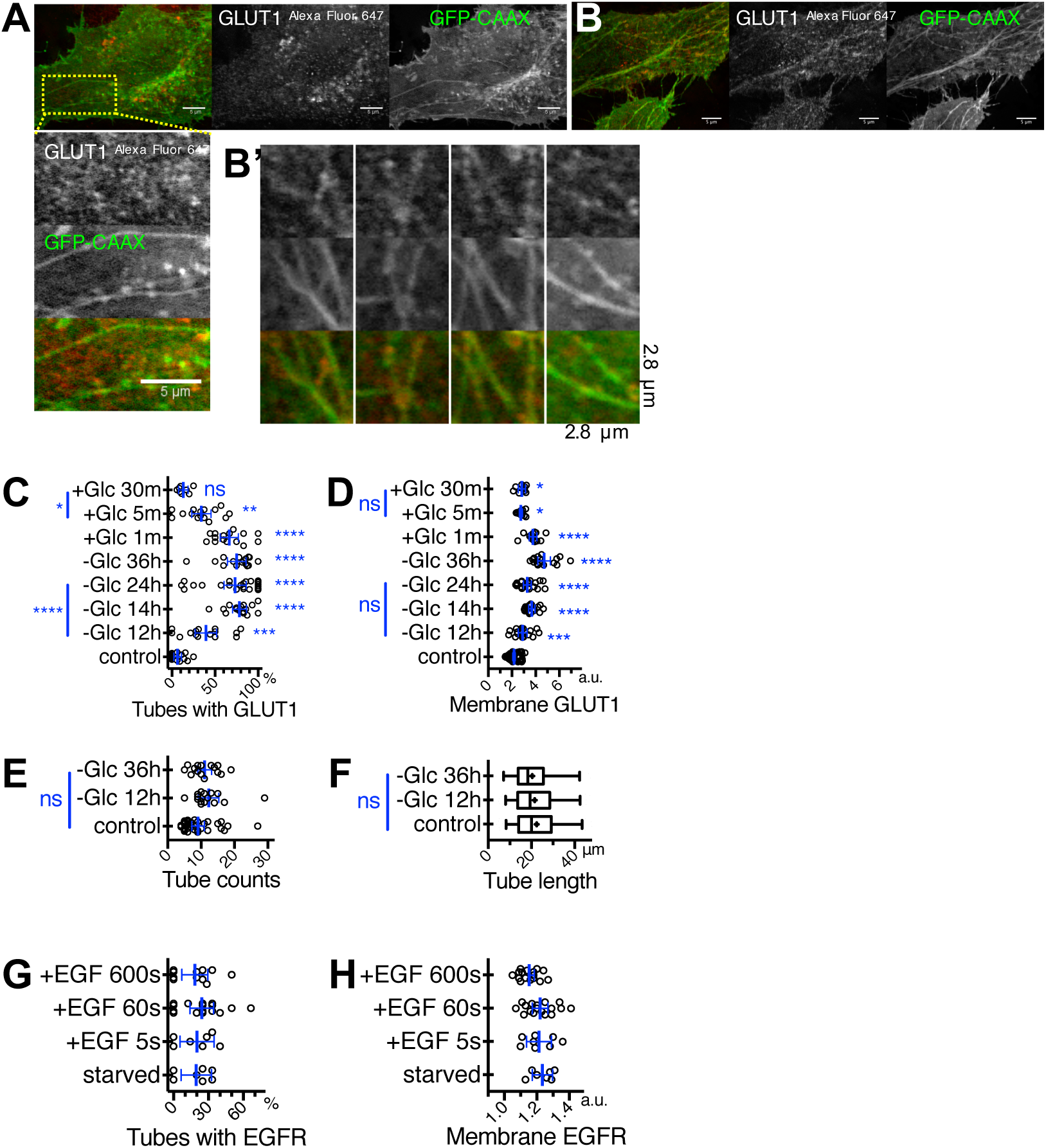
The plasma membrane tubes facilitate receptor presentation to the surface of cells. A- HeLa cells expressing GFP-CAAX were immunolabeled by antibody against GLUT1 and imaged by SDCM. Z stacks from optical sectioning were combined by wavelet-based extended depth of focus projection and presented with pseudo colors, GLUT1 in red and GFP-CAAX in green. Expanded views of yellow dash lines boxed region are displayed in the lower panels. Scale bar = 5 μm. B,B’- HeLa cells as in (A) were glucose-deprived for 12 hr, processed and presented as in (A). Four different regions are displayed in B’ in expanded view. C,G- HeLa cells expressing GFP-CAAX were treated under different conditions, fixed, immunolabeled by antibody against either GLUT1 or EGFR, and imaged by SDCM. The percentage of tubes positive for GLUT1/EGFR are presented as scatter dot plots, with mean ± 95% confidence intervals. Turkey multiple comparison test was used after one-way ANOVA to generate the p values (****, P<0.0001; ***, P<0.001; **, P<0.01; *, P<0.05; ns, not significant). Glc is the abbreviation for glucose. D,H- For cells in (C,G), the average pixel intensity of GLUT1/EGFR on the plasma membrane divided by the background are presented as scatter dot plots, and statistically analyzed as in (C,G). E,F- For cells in (C), the number of tubes in the cell or the length of tubes was counted or measured, and presented as scatter dot plots. One-way ANOVA reports the difference is not significant (ns).

Thus, we demonstrated that the PM tubes were used to facilitate GLUT1 surface presentation in changing environments. However, we could not observe similar behavior for EGFR (Figure 7G-H), a receptor tyrosine kinase, with its major role in signaling transduction.

## Discussion

Here, we reported that the plasma membrane (PM) extends tubular structures into the cytoplasm in cells. Inward tubulation of PM provides a shortcut for intercellular compartments to interact with the external environment and expedites membrane exchange between compartments.

### Previously characterized tubular membrane structures

Using colocalization analysis, we showed that these PM tubes were previously uncharacterized membrane structures. Below, we briefly summarize some of the known ones.

#### Cilia

Ciliary membrane protrudes from PM, which is topologically opposite to the PM tubes. While organized microtubule structures drive ciliary membrane formation, PM tubes can protrude without either actin filaments or microtubules. Functionally, cilia sense and transduce signaling by proteins enriched on the ciliary membrane (Ishikawa and Marshall, 2011). Our initial characterization suggested that the PM tubes expedite membrane exchange and receptor presentation. We believe future studies around PM tubes would be inspired by the existing investigations about the biogenesis and functions of cilia.

#### Tubular endoplasmic reticulum (ER)

ER is where secreted and membrane proteins are synthesized, and a major place for lipid manufacturing. ER is a continuous membrane structure with a single lumen, which consists of tubes and sheets. Rather than being pulled out by motors, studies have shown that reticulons and their interacting proteins are the structure components shaping the tubular ER (Hu et al., 2009; Voeltz et al., 2006). In yeast cells, cortical ER sheets would provide extensive interaction between ER and PM (Manford et al., 2012). However, in cells, the majority of ER sheets resides in perinuclear regions. We think that PM tubulation would provide increased opportunities for ER to interact with PM by interactions either between tubular ER and PM tubes, or between perinuclear ER sheets and extended PM tubes.

#### Post-Golgi tubes

The Golgi apparatus is where proteins are post-translationally modified and sorted for transport to their destinations. Many trafficking events happen in post-Golgi compartments. Using either GFP chimeras or a temperature sensitive form of the glycoprotein G of the vesicular stomatitis virus (VSV-G), studies have shown that there are tubular structures, which are Rab1 positive, being pulled along microtubules and detached from Golgi cisternae (Gallione and Rose, 1985; Presley et al., 1997; Russo et al., 2016). Post-Golgi tubes are one of the diverse ways that cells use to transport cargos to the cell periphery (Almeida et al., 2011; Anitei et al., 2010; Cooper et al., 1990; Feng et al., 2004; Ganley et al., 2008; Kreitzer et al., 2003; Lippincott-Schwartz et al., 1990; Pantazopoulou et al., 2014; Polishchuk et al., 2003; Raa et al., 2009; Sabharanjak et al., 2002; Sciaky et al., 1997; Sherer et al., 2003; Wagner et al., 1994; Waguri et al., 2003). The PM tubes characterized here display more rigid motion than post-Golgi tubes. We have shown that a small number of PM tubes could interact with the Golgi apparatus, and future investigation will be focused on when and how PM tubes connect with Golgi.

#### Elongated endosomes

In the 1990s, tubular endosomal network has been reported by two groups, after loading the cells with horseradish peroxidase for 30 min and observed through electron microscope (Tooze and Hollinshead, 1991, 1992), or after loading the cells with Texas red labelled transferrin for 60 min and observed through low light video recording (Hopkins et al., 1990). Both studies suggested a great continuity within endocytic pathways. Recently, when investigating the role of ER in endosome fission, it was shown that cells treated with dynasore or depleted of dynamin 2 by RNAi had Rab5 positive elongated endosomes (Rowland and Voeltz, 2012). Endosomes, formed by endocytosis (Antonescu et al., 2010; Howes et al., 2010), are membrane structures that internalize proteins, lipids, *etc*. All these elongated endosomes are topologically similar to PM tubes and have identical membrane source - the plasma membrane. However, PM tubes are Rab5a/b negative, transferrin receptor negative (data not shown), and some of them are stable through the cell cycle. It requires further investigation to compare the biogenesis of these two tubular structures. Also, there is Rab8-positive tubular structures that form after macropinocytosis and mediate membrane recycling and protrusion formation (Hattula et al., 2006; Roland et al., 2007). However, these Rab8-positive tubules are sensitive to nocodazole and cytochalasin D treatment, while PM tubes are not. Even though we have shown that the PM tubes connect with the plasma membrane, little is known as to where they connect. Eisosome-like domains in mammalian cells would be the first type of PM domains that we would investigate in the future (Moreira et al., 2009; Walther et al., 2006). Since dynamic tubes lack cholesterol, it is unlikely that PM tubes initiate from lipid rafts. On the contrary, existing PM tubes are likely to connect with cholesterol lipid rafts to stabilize and maintain themselves.

#### Mitochondrial and auto-lysosomal tubes

Tubulation of mitochondria and auto-lysosomes is critical in maintaining organelle homeostasis (Reyes et al., 2011; Swanson et al., 1987). Recent studies using *in vitro* reconstitution demonstrated that a kinesin motor protein, Kif5B, decorates and pulls the mitochondrial outer membrane, or the auto-lysosomal membrane, to form tubular structures (Du et al., 2016; Wang et al., 2015). Kif5B travels along microtubules at an average speed of 49~54 μm/min (Nishiyama et al., 2002; Schnitzer et al., 2000), which is comparable with the fast protrusion speed, providing one mechanism for PM tubulation. However, the frequent transmission-switching-like protrusion dynamics, as well as the ability to extend without polymerized microtubules, suggests a complex regulation that involves in PM tubulation. Future studies will focus on finding the molecular machines that are employed and comparing the difference between microtubule based and actin filament based dynamics. For each PM tube, transmission-switching-like protrusion suggests a smooth transition from microtubule-based to actin-filament based dynamics, which must involve additional coordination between the two distinct cytoskeleton networks.

In addition to the ones summarized above, in *Drosophila* respiratory system, tracheoles invade into insect flight muscle T-tubules to form close contacts with every mitochondrion (Peterson and Krasnow, 2015), providing an excellent example on how physiological needs are met by extravagant membrane deformation.

### The tunneling process

Virginia Woolf, an English writer in the twentieth century, used ‘*tunnelling process*’ to portray her characters’ state of mind. She brought a character’s past into the present moment through his memory. She described this method of writing as ‘*dig[ging] out beautiful caves*’, and she thought it gives ‘*humanity, humour, depth*’. Woolf’s tunnel implies the possibility of coming and going between the past and the present.

We demonstrated that the cell uses inward tubulation of PM to bring the external environment close to its intercellular compartments. Cellular organelles are like Woolf’s caves, having their glory moments, and more importantly, organelles shall connect, precisely and efficiently.

We propose to name the tubular membrane structures described here as the plasma membrane tunnels (PM tunnels), and name this phenomenon membrane tunneling.

The PM tunnels are distinct from previously reported nanotubules that exist between identical organelles or cells. Stroma-filled tubules are formed by tubulation of the inner and outer plastid membranes. They connect and transfer macromolecules between plastids (Kwok and Hanson, 2004; Natesan et al., 2005). To pass through plant cell wall, plasmodesmata are used. Within plasmodesmata, there are desmotubules connecting the ER of adjacent cells (Zambryski, 1995). Between mammalian cells, nanotubules form complex networks and transfer endosomal structures between cells (Rustom et al., 2004). Within mammalian cells, both short-lived and long-lasting mitochondrial nanotubules have been described, which are suggested to be important for cardiomyocyte function (Huang et al., 2013). To enable bacterial social behaviors, nanotubular superhighways directly exchange cytoplasmic factors between *B. subtilis* cells (Dubey and Ben-Yehuda, 2011). Although some of the nanotubules mentioned above bear the name of “tunneling nanotubules”, the emphasis of membrane tunneling phenomenon proposed here is not limited to the shape of the structure, but also the active process to achieve membrane exchange.

### Membrane tunneling is a general adaptation to compartmentalized cellular biology

Compartmentalized cellular biology reflects the principle that a eukaryotic cell uses to organize its internal landscape. Because of the compartmentalization, how to exchange material precisely and efficiently between different compartments is the utmost important matter a cell must deal with.

Previous studies have revealed that the cell uses membrane vesicle trafficking to precisely exchange material. Here, we described a second approach, membrane tunneling, which the cell uses to accelerate material exchange, and we provide evidence that the plasma membrane tunneling expedites membrane exchange and receptor presentation. Both membrane vesicle trafficking and membrane tunneling achieve material exchange, probably with the first being specialized in cargo delivery and the second in membrane exchange. Both approaches are probably a universal adaptation to compartmentalized cellular biology.

## Experimental procedures

### Reagents and Materials

Commercial antibodies were obtained from BD Biosciences (GM130, cat# 610822), Abcam (GLUT1, cat#115730) and Sino Biological (EGFR, cat# 1001-MM08), and Jackson Immuno Research (Alexa Flour 647 conjugated secondary antibodies). Latrunculin B from Calbiochem, thapsigargin from Sigma, nocodazole from Selleck, and paclitaxel from LC Laboratories. All other materials were from Sangon unless otherwise indicated. Cells were obtained from ATCC, or gifts from collaborating laboratories.

### Cell culture, transient transfection and viral transduction

HeLa cells were cultured in DMEM supplemented with 10% FBS (HyClone), 100 U/ml penicillin, 100 μg/ml streptomycin and 1x GlutaMax (Gibco). Transient transfections were performed using Lipofectamine 2000 (Invitrogen) and Opti-MEM (Gibco), with reduced reagent incubation time. Viral packaging, infection and fluorescence-activated cell sorting (FACS) were as described (Cai et al., 2008; Cai et al., 2007).

### Constructs and molecular cloning

PCR and subcloning were performed using standard methods. Most green fluorescent protein (GFP) fusions were mEmerald based. Red fluorescent protein (RFP) fusions were based on TagRFP-T, mApple or mCherry. Detailed primer sequences are available upon request. To convert ^SunTag^cyto into ^SunTag^CAAX, primers 5-CCC tctaga acctacaagctgatcctgaacg-3 and 5-gac AAGCTTa ggagagcacacacttgcagctcatgcagccggggccactctcatcaggag ggttcagctt CTC GGT CAC GGT GAAG-3 were used to obtain GB1-CAAX, and PCR products were cut by XbaI and HindIII, and ligated into ^SunTag^cyto to replace its GB1. All constructs were verified by Sanger sequencing.

### Dextran loading assay

HeLa cells were transfected with indicated constructs, or made stably expressing lines via viral transduction and FACS. The cells were plated onto glass bottom dishes coated with 10 μg/ml fibronectin (EMD Millipore), and allowed to grow for 12 hr. We incubated the cells with culture media containing 2mg/ml fluorescent dextran (Molecular Probes, lysine fixable dextran: Texas Red, 3,000 MW, D3328; Texas Red, 10,000 MW, D1863; Texas Red, 70,000 MW, D1864; Fluorescein, 500,000 MW, D7136; Fluorescein, 2,000,000 MW, D7137) for 10 sec, suck out the media and fixed the cells with 4% paraformaldehyde and 0.5% glutaraldehyde in Kreb’s S Buffer (145 mM NaCl, 5 mM KCl, 1.2 mM CaCl_2_, 1.3 mM MgCl_2_, 1.2 mM NaH_2_PO_4_, 10 mM glucose, 20 mM HEPES pH 7.4 and 0.4 M sucrose) for 10 min. After three sequential washes by PBS, the samples were immunostained (Cai et al., 2007) and imaged.

### Light microscopy and statistical analysis

Images, unless otherwise indicated, were captured using a spinning disk confocal scan head (CSU-X/M2N, Yokogawa) attached to an inverted microscope (IX-81, Olympus) and an EMCCD camera (pixel size = 16.016 μm; DU897BV, Andor) and controlled by Micro-Manager software. For most of the imaging, 150XOTIRF objective (N.A. = 1.45, airy disk radius = 210.3 nm) was used. To image zebrafish animal, 60XS objective (N.A. = 1.3, airy disk radius = 234.6 nm) was used.

The GFP and RFP were excited by 488-nm and 561-nm lasers. The emissions were separated by FF410/504/582/669-Di01 and further passed emission filters FF02-510/10 (mEmerald), FF01-590/20 (mApple), FF01-583/22 (TagRFP-T), FF01-615/20

(mCherry), respectively. Alignment of the GFP and RFP channels was roughly tuned with 100-nm fluorescent microspheres (TetraSpeck). Filipin and DHE imaging were performed as previously described (Hao et al., 2002; Mesmin et al., 2011). Kymograph analysis was based on timelapse movies as described in the figure legends. Statistical analyses were performed using Prism (Graphpad). All experiments were independently performed in triplicate, with at least 5 cells analyzed in each, resulting N>15; unless otherwise indicated. Representative images are shown.

## Acknowledgments

We acknowledge Tian-qi Chen for his intuitive questioning that leads to the initial discovery; a lot of people for inspiring suggestions in the early stage of the investigation; Hua Guo, Meng-yi Yao, Ling-hua Zhang and Wei Xu for critical comments on this manuscript; Tsinghua Cytoskeleton Club members and 2016 Cell Polarity Signaling GRC attendants for extensive discussions; Shu Mao for helping Filipin and DHE imaging; Hexige Saiyin for suggestion on the GLUT-1 antibody; Siyuan Wang and Xiaowei Zhuang for providing the mMaple3-vimentin plasmid, Junjie Hu for providing reticulon 4a, atlastin-1(K80A) and CLIMP-63 plasmids, Yang Chen and Li Yu for providing normal rat kidney epithelial cells (NRK), Xiang Zhao and Su Guo for providing the *Tg*(*EF1a*: myr-Tdtomato) transgenic zebrafish line, Jie Wang and Yangming Wang for providing the GCN4-sfGFP-GB1 plasmid, Xiao-li Zhang for providing the EGFP-Rab1a, EGFP-Rab5a/5b, EGFP-Rab7a, LC3-EGFP, EGFP-PXMP2 and LAMP1-TagRFP-T plasmids; Yujie Sun and Zhen Liu for suggestions in analyzing STORM data; Bi-Chang Chen, Eric Betzig, Fengzhu Xiong and Sean Megason for sharing high-resolution images of developing embryos; and the assistance of microscopy application specialists from Nikon Instruments and ZEISS. LC was supported by the National Natural Science Foundation of China grants (31222019 and 31271428), the Thousand Young Talents Plan and by funds from Fudan University and YZT Foundation.

## Author contributions

LFY, LZ and LC conceived and designed the experiments. LFY and LZ performed experiments. LFY, LZ and LC analyzed the data. RZ did the experiments in zebrafish. LC wrote the manuscript.

